# Comparative metagenomic analysis following treatment with vancomycin in C57BL/6 and BALB/c mice to elucidate host immunity and behavior

**DOI:** 10.1101/2020.05.07.083659

**Authors:** Pratikshya Ray, Debasmita Das, Uday Pandey, Palok Aich

## Abstract

The gut is the largest reservoir of the resident microbiota. The microbiota can affect the host behavior and immunity. While the consequence of treatment with antibiotics on the gut microbiota can be destructive but can be utilized as a tool to understand the host immunity and behavior. The magnitude of perturbation and time needed for the restoration of gut microbiota can depend on the immune bias of the host. In the current study, we therefore, observed the perturbation and restoration kinetics of gut microbiota following treatment with vancomycin and its effect on the host physiology in both Th1-(C57BL/6) and Th2-(BALB/c) biased mice. A comparative metagenomic analysis revealed that the treatment with vancomycin caused a significant decrease in the abundance of Firmicutes and Bacteroidetes phyla and an initial increase in Proteobacteria. Increase in Proteobacteria decreased with continued treatment with vancomycin to result into a significant rise in Verrucomicrobia phylum. We established the patterns of gut microbiota alteration and its effect on a) the behavior of mice, b) expression of key brain molecules and b) immunity related genes. We followed the gut microbiome restoration for a period of two months following withdrawal of treatment with vancomycin. Maximum restoration (>70%) of gut microbiota happened by the 15^th^ day of withdrawal. BALB/c mice showed a more efficient restoration of gut microbiota compared to C57BL/6 mice. The results, in general, revealed that along with the restoration of major gut microbes, important physiological and behavioral changes of both mice strains returned to the normal level.

## Introduction

Gastrointestinal tracts of human and other higher vertebrates harbor a large network of complex microorganisms, which is mostly host-specific and helps in maintaining proper homeostasis of the host (1–4). Any perturbation of gut microbes causes a disturbance in the homeostasis, which leads to various diseases (5, 6). Among various perturbing agents, antibiotics act as the most potent perturbing agent (7–9). The use of antibiotics not only destroys pathogens but also affects the diversity of beneficial commensal microbes present in the gut (9). Multiple studies reported the effects of different antibiotics on the composition and abundance of gut microbes (7, 10, 11). Reports suggested that overuse and abuse of antibiotics can lead to permanent changes in the composition of gut microbiota which lead to various metabolic disorders like obesity, diabetes-like diseases (8, 12, 13). Vancomycin is one of the potent antibiotics to perturb the gut microbiota (6). Vancomycin treatment caused a significant alteration in the composition and diversity of the commensal gut-microbiota of the host (14–16). The correlation, however, between the extent of gut microbiota perturbation with a specific dose and duration of vancomycin exposure is still poorly characterized. Upon cessation of antibiotic treatment, the restoration kinetics of these microbes and its effect on the host behavior, immunity and other physiological functions are still not adequately addressed. While the immune bias (Th1 and Th2) of the host may play an important role on the gut microbial composition, how will the treatment with vancomycin or following withdrawal of the treatment be affecting the innate mucosal immunity or behavior and gut microbial composition (abundance and diversity) are open questions. Th1- and Th2-biased mice are two different inbred strains that differ in their baseline microbiota composition (17, 18). However, how differentially the gut microbiota, of two differently immune biased mice (Th1- and Th2-), respond to the same dose of vancomycin that needs to be explored. The differential perturbation and restoration kinetics of gut microbiomes may affect the behavior and immunity of mice in a different way between Th1- and Th2-biased mice. Moreover, the abundance and diversity of certain microbes can significantly regulate the behavior of mice.

Earlier studies reported that altered gut microbiota or the introduction of a pathogen to the gut causes various behavioral changes like anxiety and depression in mice (19–21). Levels of Brain-derived neurotrophic growth factor (BDNF), corticotropin-releasing hormone (CRH) and CRH binding protein (CRHBP) change with the stress created by the variation of gut microbiota of mice (22–24). Dysbiosis of gut microbiota modulates the expression of various tight junction proteins causing changes in the permeability of the gut (25). Various SCFA and metabolites produced by the gut microbiota mainly regulate the expression of these tight junction proteins (26).

In the current study, we tried to establish the difference in the alteration pattern of the gut microbiota of Th1- and Th2-biased mice during and post vancomycin treatment. We also correlated the effect of perturbation and restoration kinetics of gut microbes with the behavior, expression of brain specific gene markers and immune profile of two strains of mice. The current results revealed a strong association between the abundance of specific gut microbes (*A. muciniphila, E.coli*, F/B ratio) with the altered behavior pattern of mice. Both perturbation and restoration kinetics of gut microbiota followed different patterns in the two strains of mice. This difference was reflected in their behavior and in the expression of various stress and immune regulatory genes.

## Materials and methods

### Animals Used in the study

All mice used in the present study were housed in a polysulfone cage, and corncob was used as bedding material. Two mice strains C57BL/6 (Th1-) and BALB/c (Th2-) of 6-8 weeks were used for the present study. Food and water were provided *ad libitum*. Animals were co-housed in a pathogen-free environment with a 12 h light-dark cycle (lights on from 7:00 am – 7:00 pm), temperature 24 ± 3°C and humidity 40-70% maintained. The guideline for animal usage was as per CPCSEA (Committee for the Purpose of Control and Supervision of Experiments on Animals, Govt. of India), and all protocols were approved by the Institute Animal Ethics Committee constituted by CPCSEA. A schema of the experimental protocol is shown in Fig. 1.

**Fig.1.**
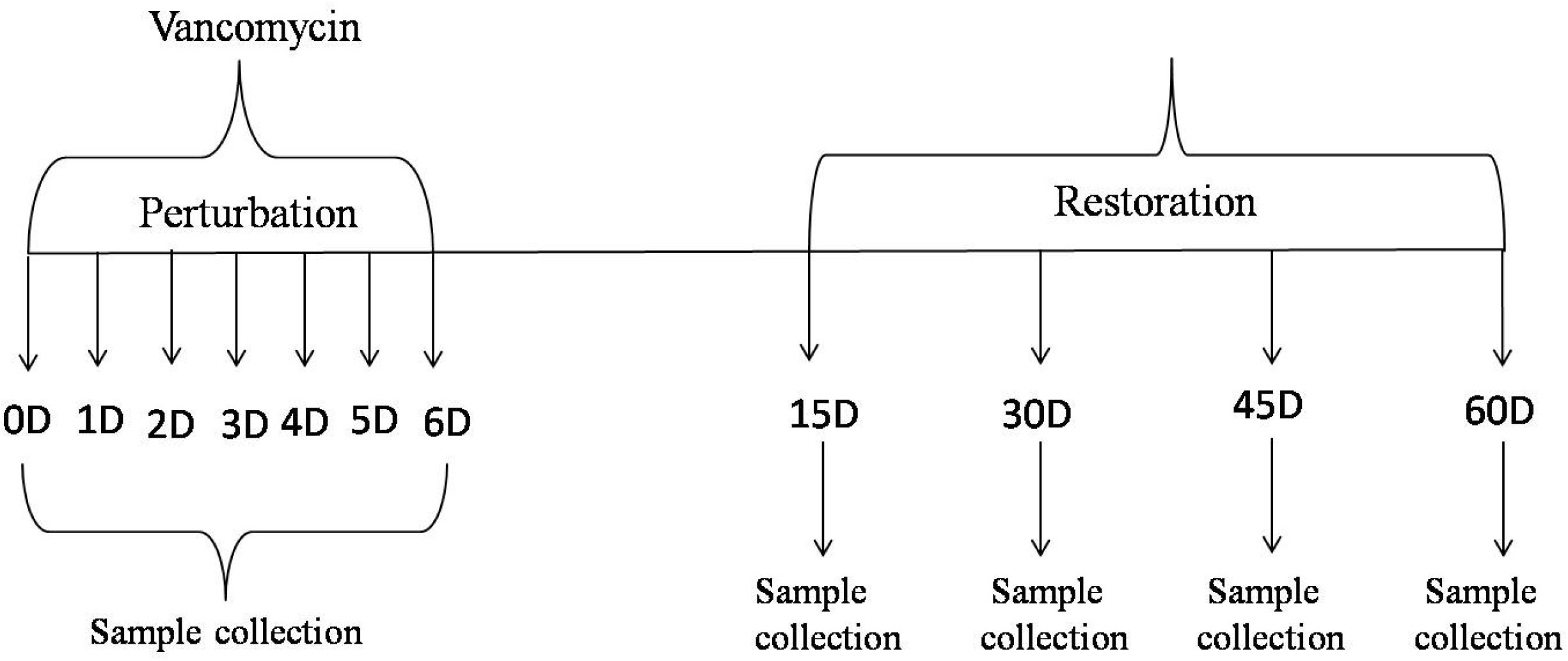
Experimental timeline of events. Experimental timeline from the study initiation day (day zero of vancomycin treatment) to the termination day (day 60 following the withdrawal of vancomycin treatment).

### Antibiotic treatment

Both Th1-(C57BL/6) and Th2-(BALB/c) biased mice were treated with vancomycin (Cat#11465492) (at 50 mg per kg of body weight) for six consecutive days (Fig.1). 0.5 ml of vancomycin was orally gavaged twice daily at a gap of 12 h. The dosage was selected as per previous reports and FDA guidelines (27, 28).

### Restoration

Following withdrawal of 6-days of treatment with vancomycin, mice were studied for 60 days. This period was termed as restoration phase. During the restoration phase, normal food and water were given *ad libitum* to mice. Mice were euthanized and various samples were collected at an interval of every 15 days of restoration, i.e., on the 15^th^, 30th, 45th and 60^th^ day (Fig.1). For the behavioral studies, separate groups of mice were used and their behaviors were observed continuously during the period of perturbation and restoration.

### Sample collection

Mice were split into two different groups: Control (untreated) and Treatment (groups that were treated with vancomycin). Mice belong to the treatment group was orally gavaged with vancomycin twice daily for 6 consecutive days. For each treated group, there was a corresponding time matched control group. Each group consisted of six mice. Mice belonged to the treated and time-matched control groups were euthanized everyday till day 6 (total of 6 time points) following treatment with vancomycin and four-time points of restoration (on 15^th^, 30^th^, 45^th^, and 60^th^ day). Mice were euthanized by using cervical dislocation method as per protocol approved by the Institutional Animal Ethics Committee. Samples were collected from Colon, brain, blood, and cecal tissues of each mouse for further analysis using the methodologies described elsewhere (29).

### RNA extraction

RNeasy mini kit (Cat# 74104, Qiagen India) was used to extract RNA from the cecal (gut) wall tissue. After sacrificing mice, the gut was washed properly and stored in RNA later for further use. During the extraction process, nearly 20-23 mg of gut tissue was churned using liquid nitrogen and 700 μl of RLT buffer was added and homogenized well. An equal volume of 70% ethanol was added and mixed well. The solution was centrifuged at 8000×g for 5 min at room temp. The clear solution containing lysate was passed through RNeasy mini column (Qiagen, Germany), which leads to the binding of RNA to the column.

The column was washed using 700 μl RW1 buffer and next with 500 μl of RPE buffer. RNA was eluted using 30 μl of nuclease-free water. RNA was quantified and the quality was checked using NanoDrop 2000 (Thermo Fisher Scientific, USA).

For RNA extraction from brain tissue, 100 mg of brain tissue was homogenized in 2 ml of TRIzol reagent. Centrifugation was done at 12000×g at 4 °C for 10 min. The fat monolayer was carefully avoided while pipetting the rest of the sample in a clean 1.5 ml MCT and 400 μL of chloroform was added to the sample. Centrifugation was again performed at 12000× g for 30 min at 4°C. RNA phase was transferred to a new MCT and 1.5 volume of 100% Ethanol was added. The sample was loaded to a spin column and HiPurA Total RNA Miniprep Purification Kit was used to extract RNA.

### cDNA preparation

cDNA was synthesized by using the AffinityScript One-Step RT-PCR Kit (Cat# 600559, Agilent, Santa Clara, US) using extracted RNA. RNA was mixed with a random 9mer primer, Taq polymerase, and NT buffer, the mixture was kept at 45°C for 30 min for the synthesis of cDNA and temperature increased to 92°C for deactivating the enzyme.

### Real-time PCR (qRT-PCR)

Real-time PCR performed in 96 well plates, using 25 ng of cDNA as template, 1 μM of each of forward (_F) and reverse (_R) primers for genes mentioned in Table 4, SYBR green master mix (Cat#A6002, Promega, Madison USA), and nuclease-free water. qRT-PCR was performed in QuantStudio 7 Real-Time PCR (Applied Biosystems, USA). All values were normalized with cycle threshold (Ct) value of GAPDH (internal control) and fold change of the desired gene was calculated with respect to control.

### Genomic DNA extraction

Fresh cecal samples from euthanized mice were collected to obtain gDNA using phenol-chloroform extraction method. 150-200 mg of the cecal sample was taken and homogenized using 1 ml of 1X PBS and centrifuged at 6700g for 10 minutes to get the pellet. The pellet was suspended in 1 ml of lysis buffer (Tris-HCl 0.1 M, EDTA 20 mM, NaCl 100 mM, 4% SDS) (pH 8) and lysed by homogenizing followed by heating at 80°C for 45 min. Lipids and proteins were removed from the supernatant by using an equal volume of phenol-chloroform extraction. This process of removing lipids and proteins was repeated till the aqueous phase became colorless. gDNA was precipitated overnight at −20°C with 3 volumes of absolute chilled ethanol. Finally, it was washed twice with 500 μl of 70% chilled ethanol and air-dried. The gDNA was dissolved in nuclease-free water and quantified using NanoDrop 2000.

### Serum collection

Mice were anaesthetized, and blood was collected by cardiac puncture. The blood sample was kept in ice for 30 mins and centrifuged at 1700×g for 15 min at 4°C, and serum was collected and stored at −80°C till further used.

### 16S rRNA sequencing (V3-V4 Metagenomics)

From cecal DNA samples, V3-V4 regions of 16S rRNA gene were amplified. For this amplification, V3F: 5’-CCTACGGGNBGCASCAG-3’ and V4R: 5’-GACTACNVGGGTATCTAATCC-3’ primer pair was used. In Illumina Miseq platform, amplicons are sequenced using paired-end (250bp×2) with a sequencing depth of 500823.1 ± 117098 reads (mean ± SD). Base composition, quality and GC content of fastq sequence were checked. More than 90% of the sequences had a Phred quality score above 30 and GC content nearly 40-60%. Conserved regions from the paired-end reads were removed. Using FLASH program, a consensus V3-V4 region sequence was constructed by removing unwanted sequences. Pre-processed reads from all the samples were pooled and clustered into Operational Taxonomic Units (OTUs) by using de novo clustering method based on their sequence similarity using UCLUST program. QIIME was used for the OTU generation and taxonomic mapping (30, 31). The representative sequence was identified for each OTU and aligned against the Greengenes core set of sequences using PyNAST program (32–35). Alignment of these representative sequences against reference chimeric data sets was done and RDP classifier against SILVA OTUs database was used for taxonomic classification.

### Calculation of Cecal index

The body weight (in gram) of individual mouse was measured and recorded. The whole cecal content from each mouse was collected and weighed. The cecal index was measured by taking the ratio of cecal content to the body weight of the respective mouse (36).

### Gut permeability test by FITC dextran

Vancomycin treated and restored mice at selected time points with the corresponding time-matched control mice were water-starved overnight. Next day FITC-dextran (Cat#F7250, Sigma-Aldrich, Missouri, US), at a concentration of 100 mg/ml, was dissolved in PBS and orally gavaged to water-starved mice. After 4 h, mice were anaesthetized by isoflurane inhalation and blood was collected by cardiac puncture. The concentration of FITC in the blood serum sample was measured by Spectrofluorometer with an excitation wavelength of 485 nm (20 nm bandwidth) and emission of 528 nm (20 nm bandwidth). The procedure was performed by following the previously described protocol (37).

### Elevated plus maze test

Elevated plus maze is commonly used for assessing anxiety levels in rodents - specifically in mice (38). It was constructed with wood, painted black and positioned 80 cm above the floor of the room. This instrument has a central platform and four crossed arms (50 cm long and 10 cm wide, each): two open and two closed arms with walls extending 30 cm above the maze floor. During the testing session, each mouse (from both untreated and antibiotics-treated groups) was placed in the center of the maze facing one of the open arms, and every animal was permitted to explore the maze for 5 mins only. During these 5 mins, the total time spent in the closed and open arms was observed. This was recorded by using a computerized video tracking system (Smart 3.0, Panlab SMART video tracking system, Harvard Apparatus). For this test total, seven mice were used (n=7).

### Forced swim test (FST)

Forced swimming test is one of the valid ways of testing despair and depression created by stress in mice model (39). A cylindrical tank (30 cm height and 20 cm diameter) was made and it was filled up to 19 cm with tap water at 24±1°C temperature. Each mouse was subjected to a 6 min of swimming session with the last five minutes considered for the data analysis. During this period, immobility was recorded by using a video camera. The mouse was considered to be immobile when it became static in the water without trying to escape. Those motions which were vital to hold its head above the water surface were not taken as immobile posture. For this test total, seven mice were used (n=7).

### Open field (OF) test

Open field test is commonly used to measure anxiety and locomotors activities in small rodents (40). The instrument was made up of wood painted in black. It is a square box illuminated by a bright light from the ceiling. Each animal was placed in the middle of the box for five mins. Its locomotor activity was measured by using a computerized video tracking system (Smart 3.0, Panlab SMART video tracking system, Harvard Apparatus). The total time spent in the periphery and center of the instrument was measured. The open field was divided by virtual lines into 16 equal squares, 12 of which constituted the peripheral zone, and the remaining 4, the central zone of the box. For this test total, seven mice were used (n=7).

### Statistical Analysis

All the graphs were plotted using GraphPad Prism version 7.0. Both ‘t’-test (to compare any 2 data sets) and ANOVA (to compare more than two datasets) were performed for statistical analysis of data as described in the text.

## Results

### Gut microbial composition of BALB/c and C57BL/6 following treatment with vancomycin

Earlier reports showed that the treatment with vancomycin could cause significant alteration of gut microbiome (15, 16). The detailed understanding of vancomycin treatment and the consequence of altered gut microbiome is yet to be established. A comparative study of time-dependent alteration pattern in the gut microbiome in two strains of mice during and post vancomycin treatment was addressed by the current group. We compared differential patterns of gut microbiota profile during perturbation and restoration period following treatment with vancomycin in BALB/c and C57BL/6 mice. Metagenomic analysis (16S rRNA) of cecal content showed a significant a) decrease in the abundance of major phyla like Firmicutes (Fig.2A and Fig.2E) and Bacteroidetes (Fig.2B and Fig.2F), and b) increase in the abundance of Proteobacteria (Fig.2C and Fig.2G) up to the fourth day of vancomycin treatment in both BALB/c and C57BL/6 mice. On day four, following treatment with vancomycin, the abundance of Proteobacteria phylum was the highest (nearly 80% in both BALB/c and C57BL/6 mice). At a later stage of vancomycin treatment (after day four), the abundance of Verrucomicrobia phylum increased significantly in C57BL/6 mice compared to BALB/c mice (41). On the sixth day following vancomycin treatment, Verrucomicrobia abundance was nearly 30% in BALB/c mice and 72% in C57BL/6 mice (Figs. 2D and 2H).

**Fig.2.**
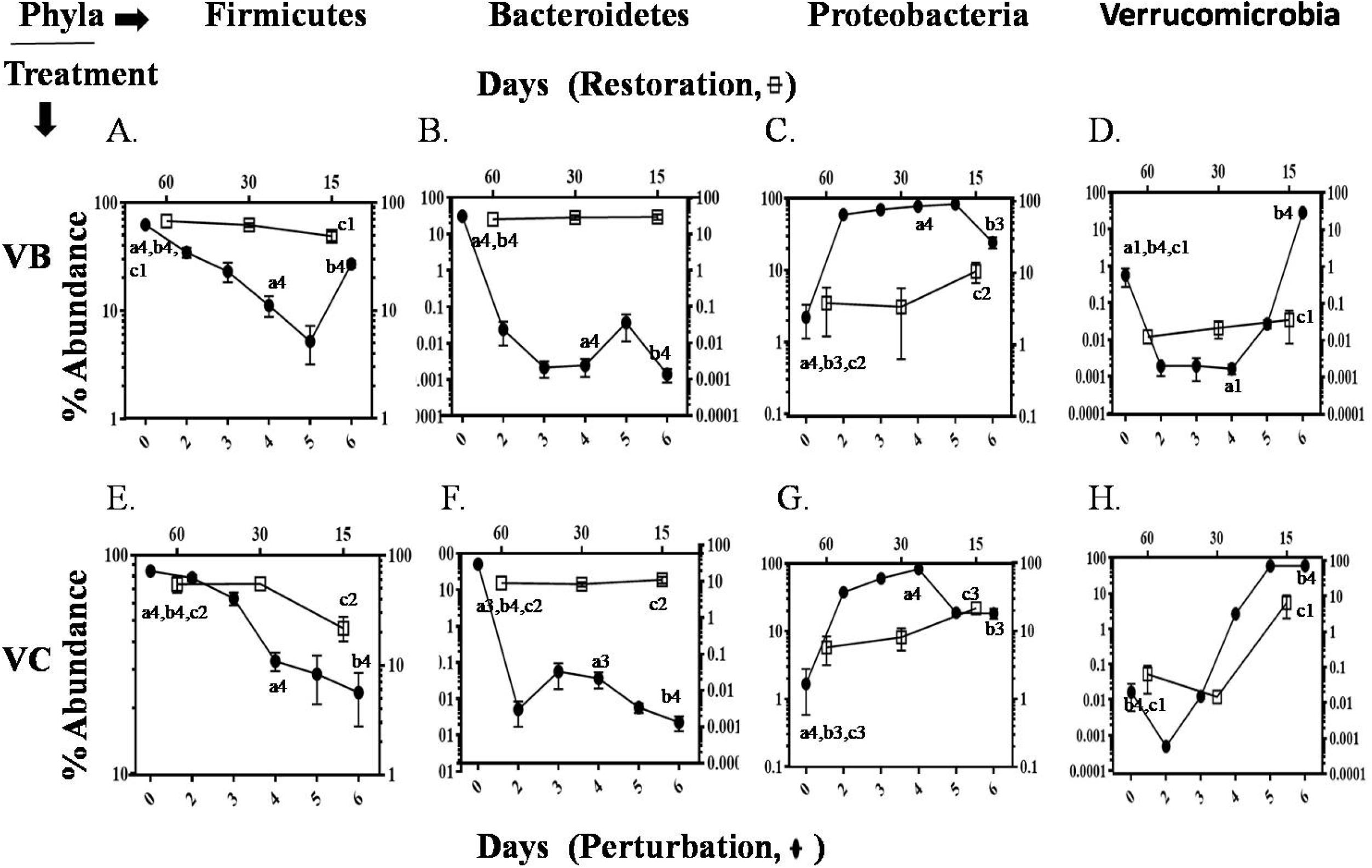
Perturbation and Restoration kinetics. Time kinetics of the major phyla (denoted by the name on top of each column of the panels) of gut microbiota following treatment with vancomycin till day 6 (perturbation) and withdrawal of vancomycin post day 6 (restoration). Time-dependent percent changes in gut microbiota abundance during vancomycin perturbation and restoration, (Top Row) in the BALB/c mice A. Firmicutes phylum, B. Bacteroidetes phylum C. Proteobacteria phylum D. Verrucomicrobia phylum and (Bottom Row) in C57BL/6 mice, E. Firmicutes phylum, F. Bacteroidetes phylum G. Proteobacteria phylum H. Verrucomicrobia phylum. The lower X-axis represents perturbation days, and the upper X-axis represents restoration days while-axis represents percentage abundance of gut microbiota. VB denotes vancomycin treated BALB/c and VC denotes vancomycin treated C57BL/6.Statistical significance changes were calculated by comparing values of the treated groups at various time points with their respective untreated groups using either two–way ANOVA or t-test, as described in preceding sections. ‘a’ showed Comparison between zero day and fourth day of perturbation; ‘b’ showed comparison between zero day and 6^th^ day of perturbation; ‘c’ showed comparison between zero day and 15^th^ day of restoration. a1, and c1 corresponds to P≤0.05; c2 corresponds to P≤0.01; a3, b3, c3 corresponds to P≤0.001; a4, b4 corresponds to P≤0.0001. Error bars shown are a standard deviation from the mean value of three replicates.

We stopped vancomycin treatment after the sixth day and left the mice to recover (termed as restoration phase). We observed the restoration of gut microbiota on the 15^th^, 30^th^ and 60^th^ day following the withdrawal of the treatment with vancomycin. The metagenomic data of cecal content during the restoration phase showed an increase in Firmicutes and Bacteroidetes phyla and a decrease in Proteobacteria and Verrucomicrobia phyla in both BALB/c and C57BL/6 mice (Figs.2 and 3). Overall, BALB/c mice showed higher efficiency in restoring the gut microbiota. The results revealed that the composition of BALB/c mice became similar to its respective control mice faster than C57BL/6 mice (Fig.3). A few selected time points were chosen to show major transitions in the gut microbial abundance and diversity during the perturbation and restoration period (Fig.3). On the 15^th^ day of restoration, in C57BL/6 mice, nearly 16% higher Proteobacteria and 18% lower Bacteroidetes phyla (Fig.3I) were observed compared to its respective time matched control group of mice (Fig. 3F). While, in BALB/c mice, only 8% higher Proteobacteria and no significant difference in Bacteroidetes phyla (Fig.3D) were found compared to its untreated control mice (Fig. 3A).

**Fig.3.**
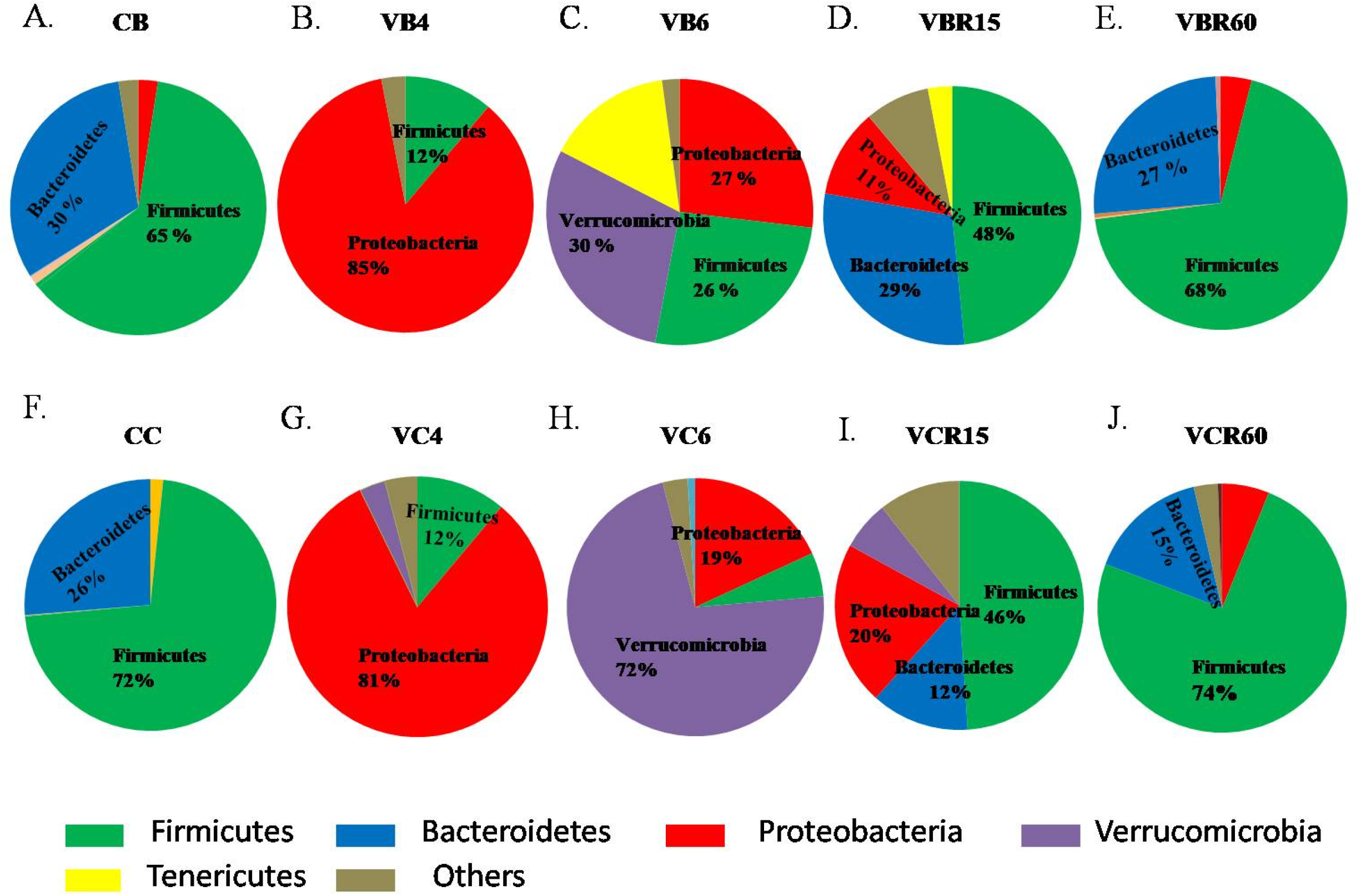
Pie chart showing the comparative changes in major phyla of gut microbiota at important time points of the experiment: Zero day (untreated mice), fourth and sixth day of vancomycin perturbation, 15th and 60th day of restoration in both BALB/c and C57BL/6 mice. Abundance of major phyla of gut microbes (Top Row) in BALB/c mice on day A. 0 (untreated control)(CB), B. 4(VB4), C. 6(VB6) following treatment with vancomycin, orD. 15(VBR15), and E. 60(VBR60)following withdrawal of vancomycin treatment, (Bottom Row) in C57BL/6 mice on day F. 0 (CC), G. 4(VC4), and H. 6(VC6) following treatment with vancomycin, or I. 15 (VCR15), and J. 60 (VCR60)following withdrawal of vancomycin treatment. Color codes of each phylum are shown at the bottom of the figure.

On the 60^th^ day of restoration, in BALB/c mice, maximum gut microbiota from all the major phyla was restored and looked almost similar to the microbiota of untreated control mice (Fig. 3). In C57BL/6 mice, Bacteroidetes and Proteobacteria phyla were not fully restored. Bacteroidetes phylum was nearly 10% lower and Proteobacteria phylum was nearly 4% higher compared to the respective untreated group of mice. The difference, in the gut microbiota level between the two strains of mice on the 60^th^ day of restoration, was significant. The restoration was more effective in BALB/c (Figs. 3A and 3E) than C57BL/6 mice (Figs. 3F and 3J) (Table 1).

**Table 1:**
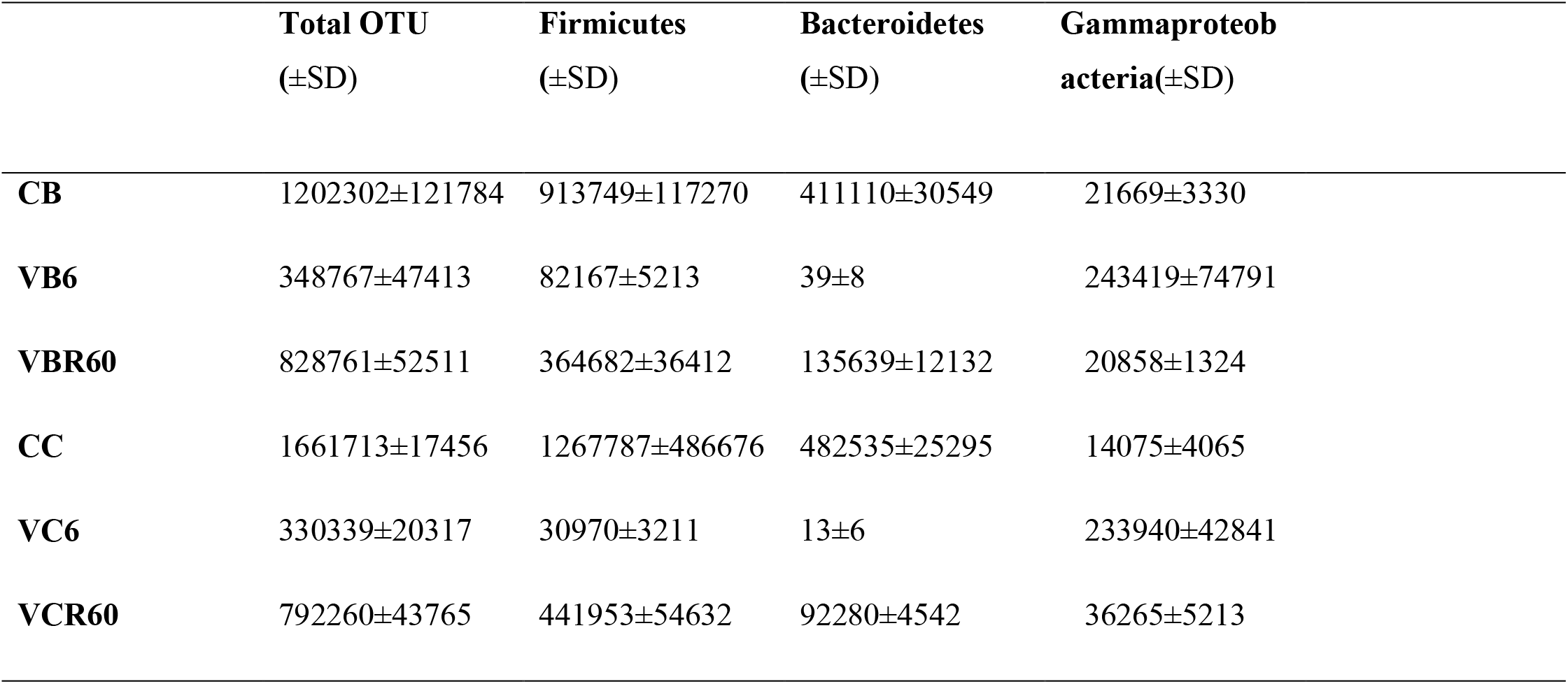
Total bacterial Operational Taxonomic Unit (OTU) abundance, as determined using metataxonomic studies by 16S rRNA analysis shown. Total bacterial OTU number, and some major groups of bacterial OTU number present in the cecal content of mice determined by metagenomic analysis (16S rRNA) in different conditions (vancomycin treated groups of mice (VB6, VC6) along with the time matched control mice (CB6, CC6) in BALB/c and C57BL/6 mice. VBR60, VCR60 correspond to 60^th^ day restored mice following cessation of vancomycin treatment).

| | **Total OTU**<br>(±SD) | **Firmicutes**<br>(±SD) | **Bacteroidetes**<br>(±SD) | **Gammaproteobacteria**(±SD) |
| --- | --- | --- | --- | --- |
| **CB** | 1202302±121784 | 913749±117270 | 411110±30549 | 21669±3330 |
| **VB6** | 348767±47413 | 82167±5213 | 39±8 | 243419±74791 |
| **VBR60** | 828761±52511 | 364682±36412 | 135639±12132 | 20858±1324 |
| **CC** | 1661713±17456 | 1267787±486676 | 482535±25295 | 14075±4065 |
| **VC6** | 330339±20317 | 30970±3211 | 13±6 | 233940±42841 |
| **VCR60** | 792260±43765 | 441953±54632 | 92280±4542 | 36265±5213 |

In addition, the diversity of gut microbiota decreased during vancomycin treatment for both strains of mice. Shannon diversity index (H) at the phylum level was found to be the lowest on day four following vancomycin treatment in both BALB/c and C57BL/6 mice. During the restoration period, H value increased and became similar to that of the untreated mice (Table 2).

**Table 2:** Shannon diversity index (H) of BALB/c and C57BL/6 mice at phylum level. Diversity index was calculated at major time points of the perturbation and restoration period of gut microbiota.

| **Perturbation Days** | **H of BALB/c** | **H of C57BL/6** |
| --- | --- | --- |
| 0 | 0.97±0.05 | 0.8±0.02 |
| 4 | 0.37±0.04 | 0.4±0.07 |
| 6 | 0.9±0.09 | 0.78±0.03 |
| **Restoration Days** |  |  |
| 15 | 1.1±0.08 | 0.9±0.07 |
| 30 | 0.9±0.05 | 0.8±0.02 |
| 60 | 0.8±0.06 | 0.8±0.04 |

### Altered gut microbiota increased anxiety and depressive-like behavior in mice

Gut microbiota has major effects on the behavior of mice (42). Elevated plus maze (EPM), Open field (OF) and Forced swimming test (FST) techniques were widely used to study behavioral changes like anxiety and depression in mice (43–46). EPM test, during vancomycin treatment, showed that both BALB/c and C57BL/6 mice stayed longer in the closed than open arms (Figs. 4IA and 4IB) compared to its respective untreated group of mice. The still image of video captured trajectory, or the paths traversed by the mice in EPM was shown in Fig.4II. The behavior in EPM showed a higher level of anxiety in mice and it increased continuously from day zero to day six following treatment with vancomycin in BALB/C mice. In C57BL/6 mice, anxiety level increased from day zero to day four following vancomycin treatment but after day four, the anxiety level decreased. On day six, following vancomycin treatment, C57BL/6 mice spent less time in the closed arm compared to its day four. During the restoration period, anxiety level decreased and came to normal in both BALB/c and C57BL/6 mice. Mice spent less time in closed arms during the restoration period than the perturbation period. On the 15^th^ day of restoration following cessation of vancomycin treatment, BALB/c mice showed less anxiety than C57BL/6 mice. On the 60^th^ day of restoration, all the vancomycin treated groups of mice behaved nearly similar to their respective untreated group of mice (Figs. 4IA, and 4IB).

**Fig.4I.**
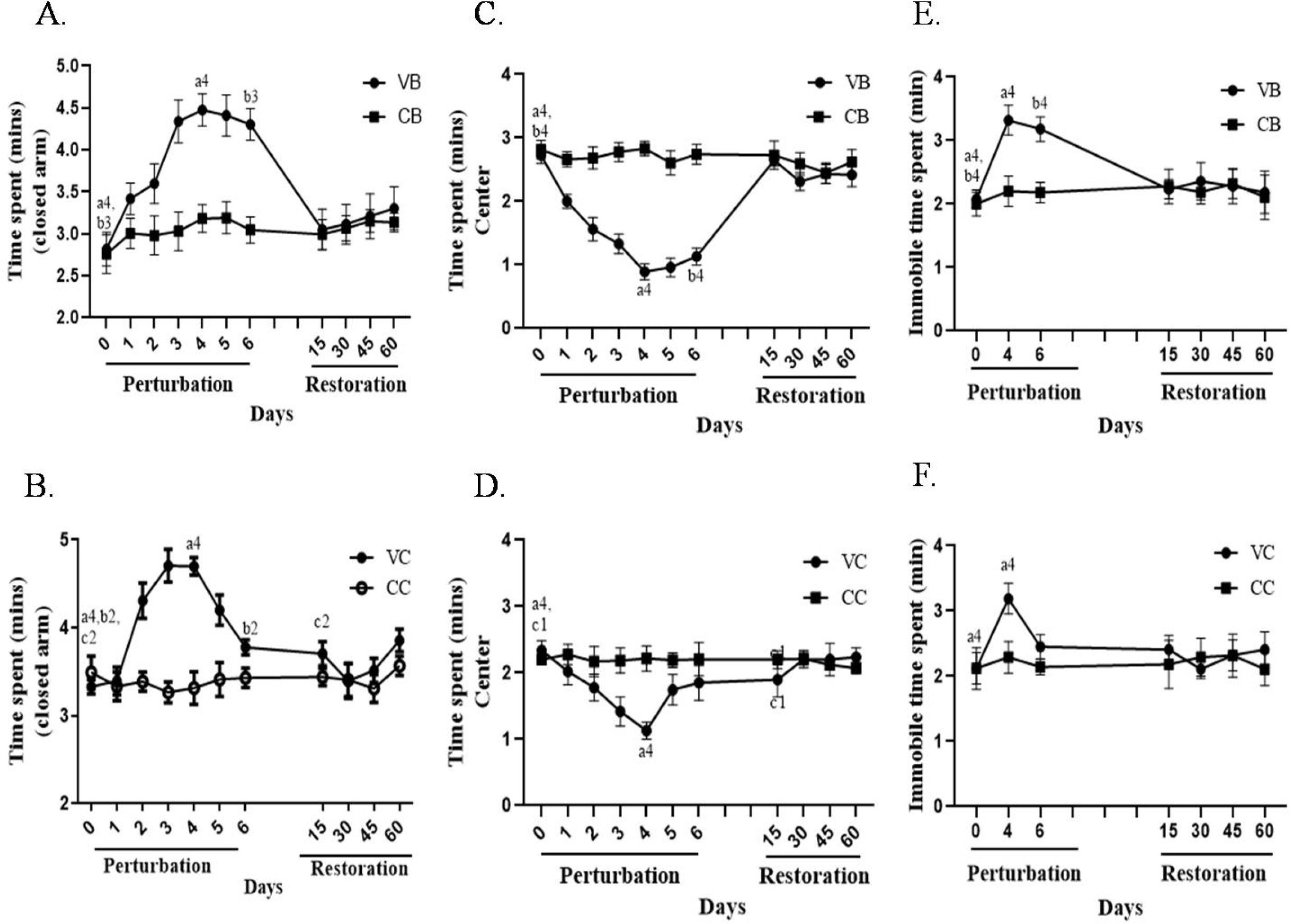
Detection of anxiety level in BALB/c and C57BL/6 mice through Elevated Plus Maze (EPM) test, open-field (OFT) and free-swimming test (FST). Elevated plus-maze data showing time spent in the closed arms (in minutes) for vancomycin treated A. BALB/c (VB) and untreated control mice (CB), or B. C57BL/6 (VC) and untreated control mice (CC) during various time points of gut microbiota perturbation (following treatment with vancomycin)and restoration (withdrawal of vancomycin treatment). Open Field Test data showing time spent in the center (in minutes) for vancomycin-treated C. BALB/c (VB) and untreated control mice (CB), or D. C57BL/6 (VC) and untreated control mice (CC) during various time points of gut microbiota perturbation and restoration. Forced swimming test data showing Immobility time spent (in minutes) for vancomycin-treated E. BALB/c (VB) and untreated control mice (CB), or F. C57BL/6 (VC) and untreated control mice (CC) during various time points of gut microbiota perturbation and restoration. (Statistical significance changes were calculated by comparing values of the treated groups at various time points with their respective untreated groups through two-way ANOVA and t-test. ‘a’ showed Comparison between zero day and fourth day of perturbation; ‘b’ showed comparison between zero day and 6^th^ day of perturbation; ‘c’ showed comparison between zero day and 15^th^ day of restoration.) cl corresponds to P≤0.05; b2,c2 corresponds to P≤0.01; b3 corresponds to P≤0.001; a4,c4 corresponds to P≤0.0001. Error bars shown are a standard deviation from the mean value of seven replicates (n=7).

**Fig.4II.**
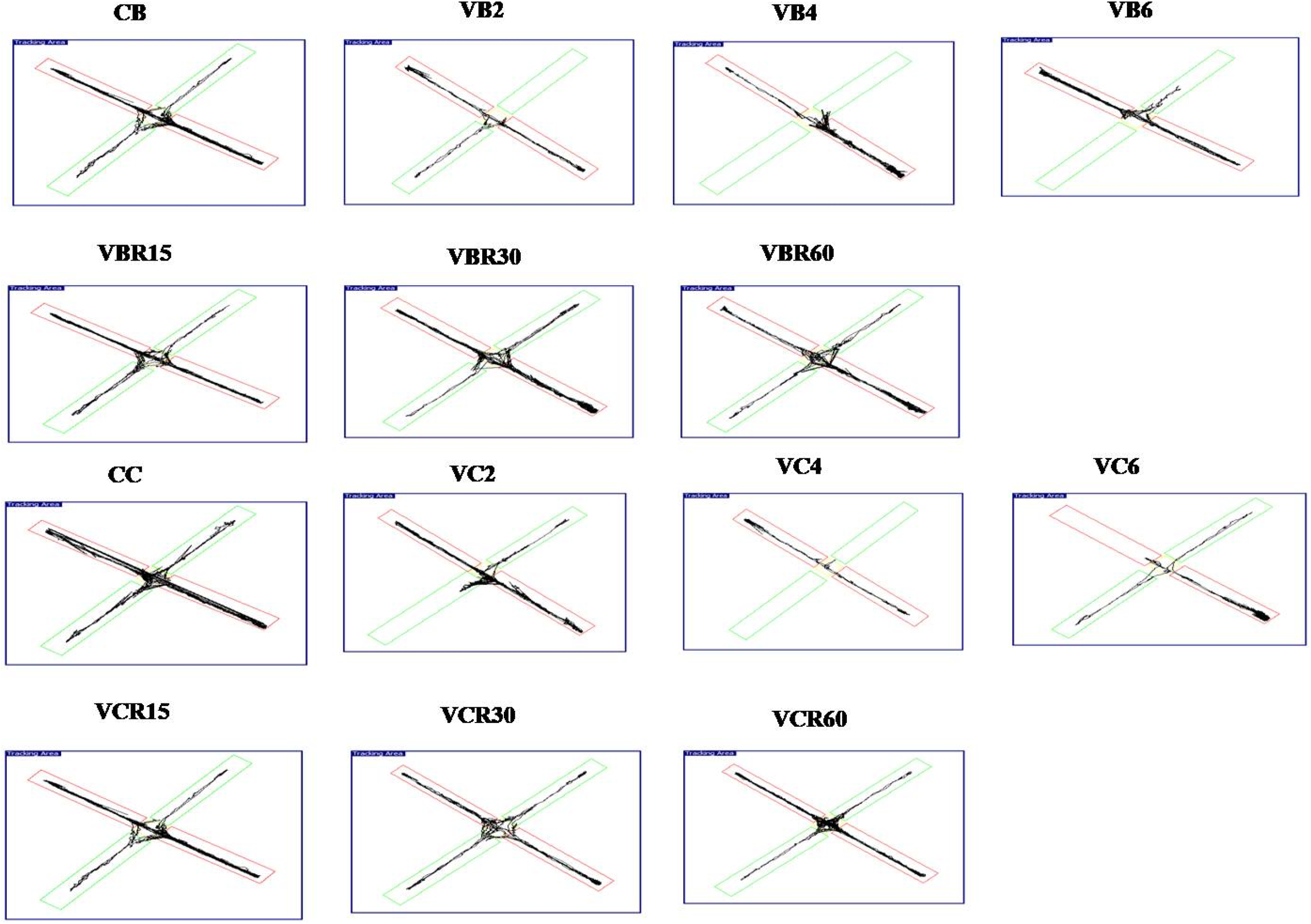
Locomotor activities by the Elevated Plus Maze test. Image depicts the trajectory or the paths that the mice traversed to represent the locomotor activities at different location of the Elevated plus maze instrument during vancomycin treatment and restoration phase of both BALB/c and C57BL/6 mice. Tracking areas are measured by Smart 3.0, Panlab SMART video tracking system, Harvard Apparatus. Untreated control BALB/c (CB) and C57BL/6 mice(CC); day 2, 4, and 6 days following vancomycin treatment in BALB/c mice (VB2, VB4, VB6), and C57BL/6 (VC2, VC4, VC6); day 15, 30, 45, 60 days following withdrawal of vancomycin treatment in BALB/c (VBR15, VBR30, VBR45, VBR60) and C57BL/6 (VCR15, VCR30, VCR45, VCR60).

Open field (OF) test showed similar results like elevated plus-maze for vancomycin treated mice. A higher level of anxiety was found in vancomycin treated mice than control mice. The results from the OF test showed that during vancomycin treatment, mice spent less time in the center than control mice (Figs. 4IC and 4ID). The still image of video captured trajectory, or the paths traversed by the mice in OFT is shown in Fig.4III. Up to the day four following vancomycin treatment, both BALB/c and C57BL/6 mice showed an increase in anxiety-like behavior (less time spent in the center). However, from day four to six following vancomycin treatment, C57BL/6 mice showed significantly less anxiety-like behavior (more time spent in the center) compared to BALB/c mice.

**Fig.4III.**
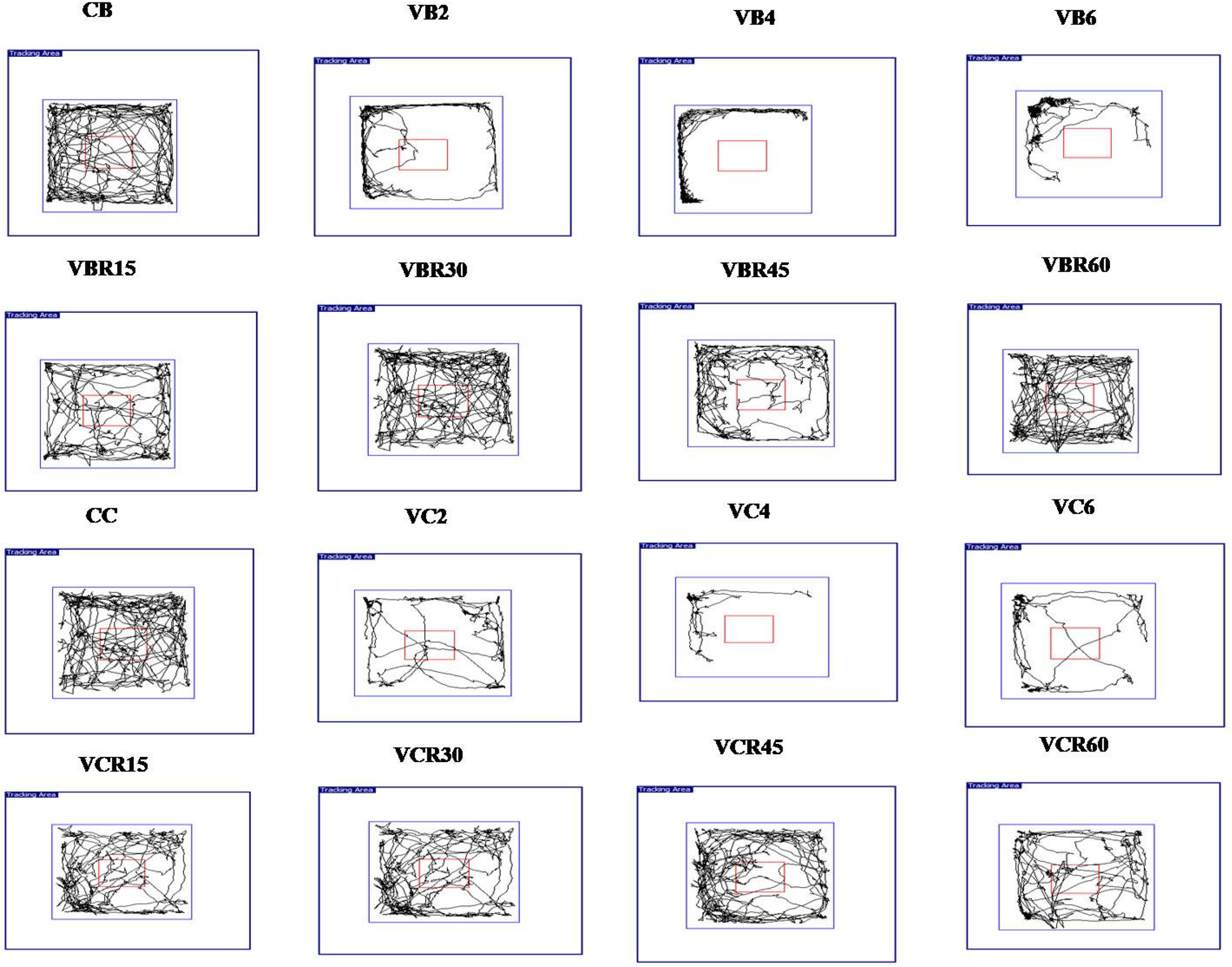
Locomotor activities by the Open Field Test. Image depicts the trajectory or the paths that the mice traversed to represent the locomotor activities at different locations of the open field instrument during vancomycin treatment and restoration phase of both BALB/c and C57BL/6 mice. Images were taken and tracking areas are measured by Smart 3.0, Panlab SMART video tracking system, Harvard Apparatus. Untreated control BALB/c (CB) and C57BL/6 mice(CC); day 2, 4, and 6 days following vancomycin treatment in BALB/c mice (VB2, VB4, VB6), and C57BL/6 (VC2, VC4, VC6); day 15, 30, 45, 60 days following withdrawal of vancomycin treatment in BALB/c (VBR15, VBR30, VBR45, VBR60) and C57BL/6 (VCR15, VCR30, VCR45, VCR60).

During the restoration period, vancomycin treated both BALB/c and C57BL/6 mice started to spend more time in the center compared to their perturbation period, which showed a decrease in the anxiety level. While for BALB/c mice there was no significant difference in the center time was found, C57BL/6 mice showed a significant difference between 15th day restoration and time matched control. On the 60^th^ day of restoration both vancomycin treated BALB/c and C57BL/6 mice behave nearly similar to their respective control group of mice (Figs. 4IC and 4ID).

In Forced swimming test (FST), it was found that the vancomycin treated group of mice became immobile most of the time in the water without trying to escape, which showed a higher level of depression in these mice compared to the control mice (Figs. 4IE and 4IF). This depressive behavior was highest on the day four following vancomycin treatment in both BALB/c and C57BL/6 strains. While on the day six following vancomycin treatment, a significantly less depressive behavior was found in C57BL/6 mice than BALB/c mice. On the 15^th^ day of restoration, vancomycin treated BALB/c and C57BL/6 mice behaved nearly similar to control mice that showed a significant decrease in their depression level within 15 days of restoration.

In summary, gut microbiota perturbation following vancomycin treatment led to significant behavioral changes as examined by the open field, elevated plus maze and forced swim tests. The recovery in the behavioral changes is associated with the time-dependent restoration of gut microbiota profile.

### Antibiotic treatment changed BDNF and CRH levels in the mouse brain

Change in behavior is an indication of changes in brain function. As described before, there are a few signature molecules, brain-derived neurotrophic factor (BDNF) and corticotropin-releasing hormone (CRH), whose expression levels speak volumes (47, 48). BDNF is necessary for the maintenance of neuronal circuit formation and its level of expression in the brain is associated with depression and anxiety of the host (47). Gut microbiota has a significant role in regulating BDNF expression (46, 49). We studied mRNA level expression of BDNF from the hippocampus of both antibiotic perturbed and restored mice. It was found that the BDNF level was decreased in both vancomycin treated BALB/c and C57BL/6 mice (Figs. 5A and 5B). Up to day four following vancomycin treatment, both BALB/c and C57BL/6 mice showed a decrease in BDNF expression. On the day six following vancomycin treatment, this reduction in the expression of BDNF was lower in C57BL/6 mice compared to BALB/c mice. During the restoration process, within the 15 days of withdrawal of vancomycin, the BDNF level came to nearly normal level in the brain of vancomycin treated mice.

**Fig.5.**
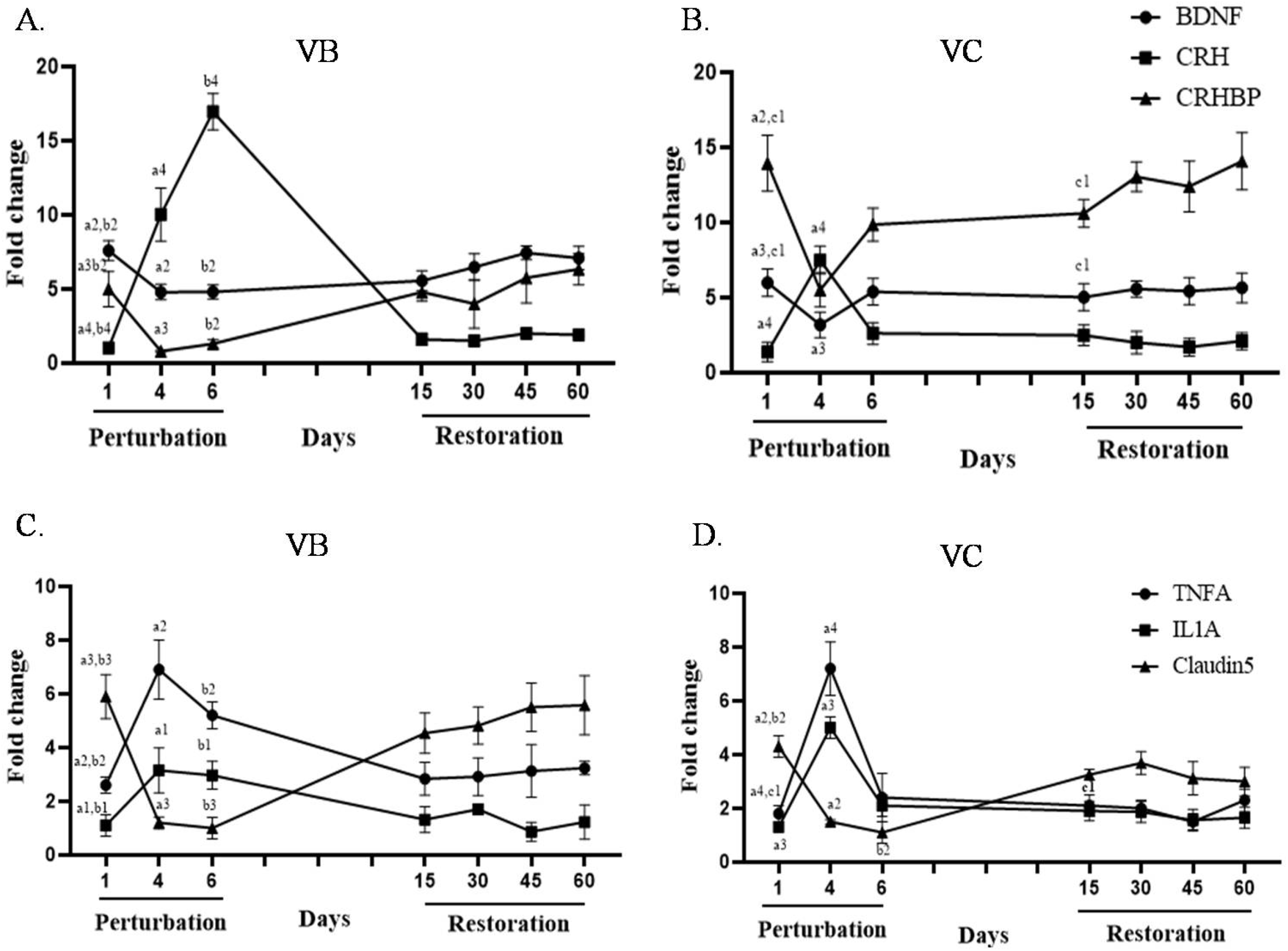
Transcriptional profile of different genes in the brain of mice. Kinetics of expression (by qRT-PCR) of various stress-related and inflammatory genes in the brain of the mice during vancomycin perturbation and restoration period. BDNF, CRH, CRHBP gene expression at mRNA level in A. BALB/c (VB) and B. C57BL/6 (VC) mice. Immune (tnfα, il1a) and tight junction genes (claudin5) expression at mRNA level in the brain of C. BALB/c (VB) and D. C57BL/6 (VC) mice. (Statistical significance changes were calculated by comparing values of the treated groups at various time points with their respective untreated groups through two-way ANOVA and t-test. ‘a’ showed Comparison between zero day and fourth day of perturbation; ‘b’ showed comparison between zero day and 6^th^ day of perturbation; ‘c’ showed comparison between zero day and 15^th^ day of restoration.) a1, b1 and c1 corresponds to P≤0.05; a2,b2 corresponds to P≤0.01; a3,b3 corresponds to P≤0.001; a4,b4 corresponds to P≤0.0001. Error bars shown are a standard deviation from the mean value of six replicates.

Like BDNF, gut microbiota also modulates CRH in the hypothalamus of the brain and regulates stress response in the host (50). We tested the mRNA level expression of CRH and CRHBP in the vancomycin treated mice brain hypothalamus and found a higher level of CRH and lower level of CRHBP compared to control mice (Figs. 5A and 5B). During the restoration process, both CRH and CRHBP came to the nearly normal level within 15 days of cessation of vancomycin. On the 15^th^ day of restoration, BALB/c mice showed higher similarity in the expression of BDNF, CRH, and CRHBP with their control groups, while C57BL/6 mice showed less similarity with their control group.

### The inflammatory response changed in the gut and brain with antibiotic treatment

Gut microbiota regulates the immune response and inflammatory state of the brain and its perturbation can cause cytokine-induced depression in the host (42, 51). In the current study, we checked the expression of select cytokines like tnfα, il1a and il10 genes at mRNA level in mice brain by qRT PCR. During vancomycin treatment, it was found that the levels of both tnfα and il1a increased significantly in the brain of BALB/c and C57BL/6 mice (Figs. 5C and 5D). Their expression was highest on the day four following vancomycin treatment. No significant changes were found in the expression of il10 in the brain of mice during vancomycin treatment (data not shown to avoid clutter).

Tight junction protein expression in the brain maintains the integrity of the blood-brain barrier (BBB). BBB of germ-free mice is more permeable than their SPF counterparts (26), which showed the significance of gut microbiota in maintaining healthy BBB. In this study, we checked the expression of claudin5 at the mRNA level in the brain of both BALB/c and C57BL/6 mice following antibiotic treatment. A significant reduction was observed in the expression of claudin5 gene at the mRNA level in the vancomycin treated mice brain compared to its time-matched untreated group of mice during the perturbation period (Figs. 5C and 5D). On the 15^th^ day of restoration, we observed an increase in claudin5 expression in the brain and became similar to the control group of mice.

We also studied the changes in the cytokines at the mRNA level in the gut during perturbation and restoration of gut microbes. We found a significant increase in pro-inflammatory cytokines gene expressions like tnfα and il1a (Figs. 6A and 6B) and a decrease in anti-inflammatory cytokine-like il10 (Fig. 6C) in the gut following vancomycin perturbation. On the day four following vancomycin treatment, expression levels of tnfα and il1a were the highest, and expression of il10 was the lowest in both BALB/c and C57BL/6 mice with respect to its time matched control value. Within 15 days of restoration, expression of all the cytokines in the gut became similar to their respective control mice in both BALB/c and C57BL/6 mice.

**Fig.6.**
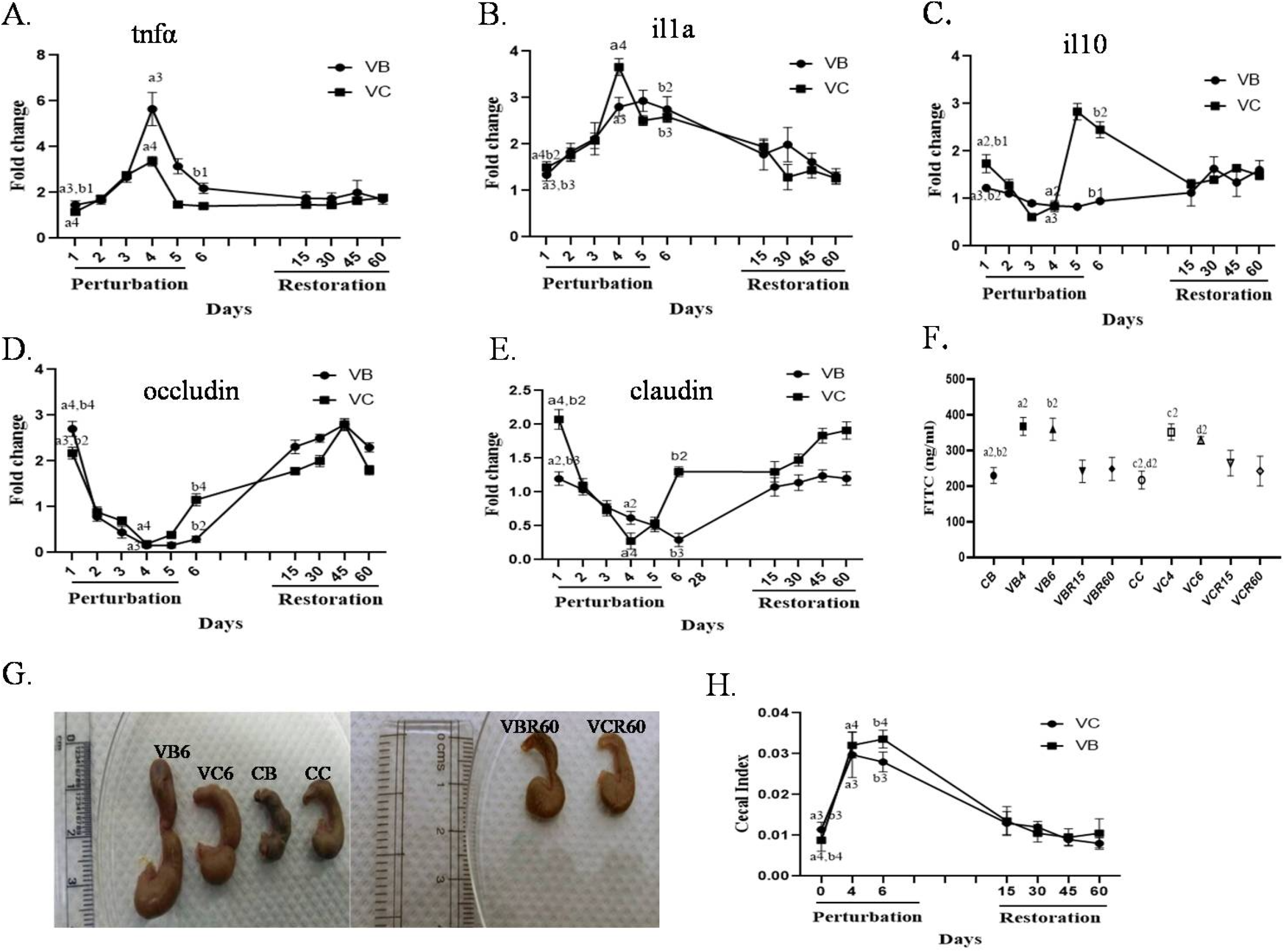
Transcriptional profile (by qRT-PCR) of various immune genes and tight junction genes in the gut tissue of mice during vancomycin perturbation and restoration period. Kinetics of expression of various immune genes at mRNA level in the colon of vancomycin treated BALB/c (VB) and C57BL/6 (VC) mice, A. tnfα, B. il1a C. il10, and tight junction genes D. occludin and E. Claudin1.F. FITC dextran concentration in serum at various time points of perturbation and restoration period. G. Representative images of various sizes of cecum from both BALB/c and C57BL/6 mice. (Cecum of day 6 following vancomycin treatment in BALB/c (VB6) and C57BL/6 (VC6), day 60 following the withdrawal of vancomycin treatment in BALB/c (VBR60) and C57BL/6 (VCR60), untreated control mice C57BL/6 (CC) and BALB/c (CB)) H. kinetics of the cecal index during gut microbiota perturbation and restoration period. (Statistical significance changes were calculated by comparing values of the treated groups at various time points with their respective untreated groups through two-way ANOVA and t-test. ‘a’ showed Comparison between zero day and fourth day of perturbation; ‘b’ showed comparison between zero day and 6^th^ day of perturbation; ‘c’ showed comparison between zero day and 15^th^ day of restoration.) b1 and c1 corresponds to P≤0.05; a2,b2,c2,d2 corresponds to P≤0.01; a3,b3 corresponds to P≤0.001; a4,b4 corresponds to P≤0.0001. Error bars shown are a standard deviation from the mean value of six replicates.

### Antibiotic treatment increased gut permeability by modulating the expression of tight junction protein

The tight junction proteins like Occludin and Claudin regulate the integrity of the gut (52). Alteration of gut microbiota could compromise the expression of tight junction proteins and might lead to inflammation (53). In the current study, we found a lower expression of occludin (Fig. 6D) and claudin1 (Fig. 6E) at mRNA level in colon tissue during vancomycin treatment in both BALB/C and C57BL/6 mice, which might be a reason for the increased permeability of the gut. During the restoration process, their expression increased and became similar to the control mice within 15 days of the restoration period. For further confirmation of gut permeability, FITC conjugated dextran was gavaged to the day 4 mice following vancomycin treatment and day 60 mice of restoration group following cessation of antibiotic treatment. FITC dextran concentration was found to be significantly higher on the 4^th^ day following vancomycin treatment in both Th1- and Th2-biased mice (367±25 ng/ml in BALB/c and 350±23 ng/ml in C57BL/6) (Fig. 6F). During the restoration period, FITC concentration in the serum came to the normal level.

Gut microbiota abundance and composition regulate the cecal size of mice. The cecum is known to be a better representation than the fecal sample for understanding intestinal microbiota profile and cecum size changes during dysbiosis of gut microbiota (54, 55). A Large cecum was observed in vancomycin treated mice than the untreated group of mice (Fig. 6G). The cecal index was calculated and found to be increased continuously during the perturbation period in both BALB/c and C57BL/6 mice (Fig. 6H). During the restoration process, the cecum size decreased and came to the normal level within fifteen days following cessation of vancomycin treatment.

## Discussion

Vancomycin perturbation caused a significant alteration in all major phyla of gut microbes (14–16). In the current study, we found that the perturbation of gut microbiota caused by vancomycin and its successive restoration kinetics follow a pattern. We observed specific patterns of changes in the major phyla of gut microbes during vancomycin perturbation and restoration. These specific alteration patterns of gut microbes affected the behavior and physiology of the host significantly. We found an important correlation between certain increased (Proteobacteria, Verrucomicrobia) and decreased (Firmicutes, Bacteroidetes) gut microbes with the anxiety and depressive behavior of mice. The patterns of changes, for perturbation and restoration kinetics of gut microbiome, observed between BALB/c and C57BL/6 at the same dose of vancomycin are different. Up to day four following vancomycin treatment, we found similar extent of increase in Proteobacteria and a decrease in Firmicutes and Bacteroidetes phyla in the gut of both BALB/c and C57BL/6 mice. These similar changes in the major phyla of gut microbes had nearly the same type of impact on the physiology of both types of immune biased mice. However, after day four, a significant biased increase of Verrucomicrobia phylum in C57BL/6 mice caused differential behavioral and immunological changes between two strains of mice. We confirmed the presence of *Akkermansia muciniphila* of Verrucomicrobia phylum on the day six following vancomycin treatment in C57BL/6 mice (41).

Earlier studies established that the restoration of gut microbiota was not complete following perturbation with antibiotics. All the species of gut microbiota present before the antibiotic treatment were not recovered during the restoration period (9, 56). The current study showed that restoration of most of the gut microbiota (>70%) happened within 15 days following cessation of vancomycin treatment. On the sixty day of restoration maximum part of gut microbes became restored but their abundance and composition were not exactly similar to the untreated mice (Fig.3). Similar to the perturbation kinetics, restoration kinetics of gut microbes also varied between BALB/c and C57BL/6 mice. In BALB/c mice, restoration efficiency of certain major phyla (Proteobacteria, Firmicutes) of gut microbes was higher than C57BL/6 mice. Major changes in the gut microbiota happened on the 4^th^ and 6^th^ day of perturbation and 15^th^ day of restoration. We, therefore, mainly focused on the changes happening in the host physiology and behavior of mice on above time points.

It was reported that alteration in the composition and diversity of gut microbes caused changes in the behavior of mice in EPM, FST and OF test (26). It was also reported that colonization of germ free (GF) mice with *Bifidobacterium infantis* normalized the stress level in mice. Mono-colonization with *Escherichia coli*, however, induced even higher stress level in GF mice (43, 57). Earlier reports further suggested that different bacteria of Firmicutes phylum helped in reducing anxiety and depression like behavior in mice (58). *Akkermansia muciniphila* bacteria caused a reduction in anxiety behavior in mice (59). Reports showed that increased Firmicutes to Bacteroidetes ratio (F/B ratio) in the gut caused hypertension and anxiety-like behavior of the host (60, 61). In the current study, we observed a high-level of correlation between relative patterns of changes in gut microbiota and behavior of mice. During vancomycin treatment, up to day four, increase in pathogenic bacteria like *Escherichia coli* and decrease in beneficial bacteria caused higher anxiety and depressive behavior in both Th1- and Th2-biased mice in EPM, OF and FST tests. After day four, Verrucomicrobia phylum replaced the Proteobacteria phylum in C57BL/6 mice which caused lower anxiety and depressive behavior in C5BL/6 mice compared to BALB/c mice. On the day four following vancomycin treatment, the dominance of single phylum (Proteobacteria) caused a decrease in diversity of gut microbes and on the day six, the appearance of Verrucomicrobia and other microbes caused the increase in diversity of gut microbes. This alteration in the diversity of gut microbes was reflected in the behavior of mice. The F/B ratio increased during vancomycin treatment and decreased during the restoration period (Table 3). This increase in F/B ratio might be associated with higher anxiety behavior of mice during vancomycin treatment. During the restoration period (on the 15^th^ day), with the more efficient restoration of gut microbiota in BALB/c mice caused less anxiety and depressive behavior compared to C57BL/6 mice.

**Table 3:**
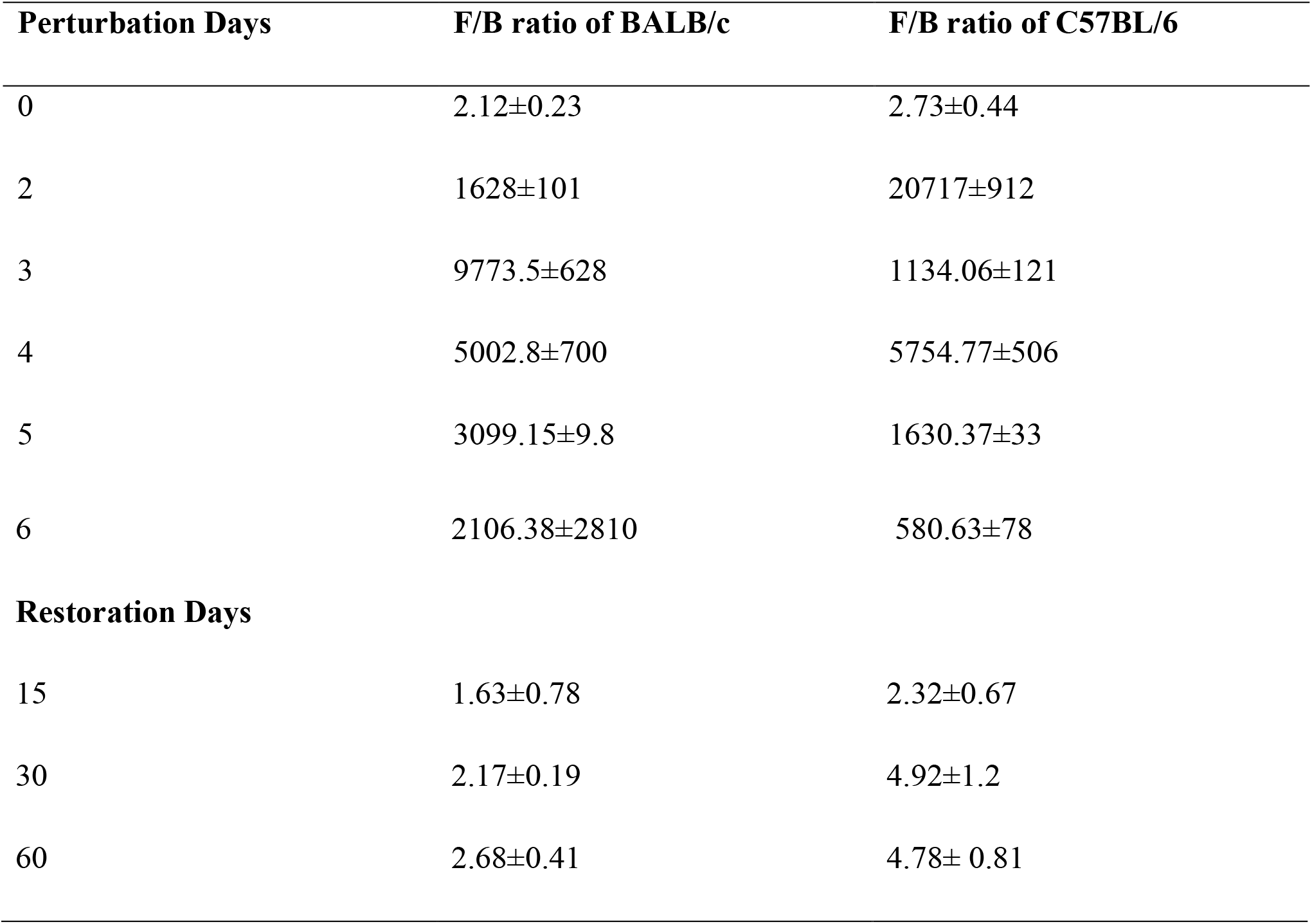
Comparison of Firmicutes to Bacteroidetes ratio (F/B) at different time points of BALB/c and C57BL/6 mice.

| **Perturbation Days** | **F/B ratio of BALB/c** | **F/B ratio of C57BL/6** |
| --- | --- | --- |
| 0 | 2.12±0.23 | 2.73±0.44 |
| 2 | 1628±101 | 20717±912 |
| 3 | 9773.5±628 | 1134.06±121 |
| 4 | 5002.8±700 | 5754.77±506 |
| 5 | 3099.15±9.8 | 1630.37±33 |
| 6 | 2106.38±2810 | 580.63±78 |
| **Restoration Days** |  |  |
| 15 | 1.63±0.78 | 2.32±0.67 |
| 30 | 2.17±0.19 | 4.92±1.2 |
| 60 | 2.68±0.41 | 4.78± 0.81 |

**Table 4:**
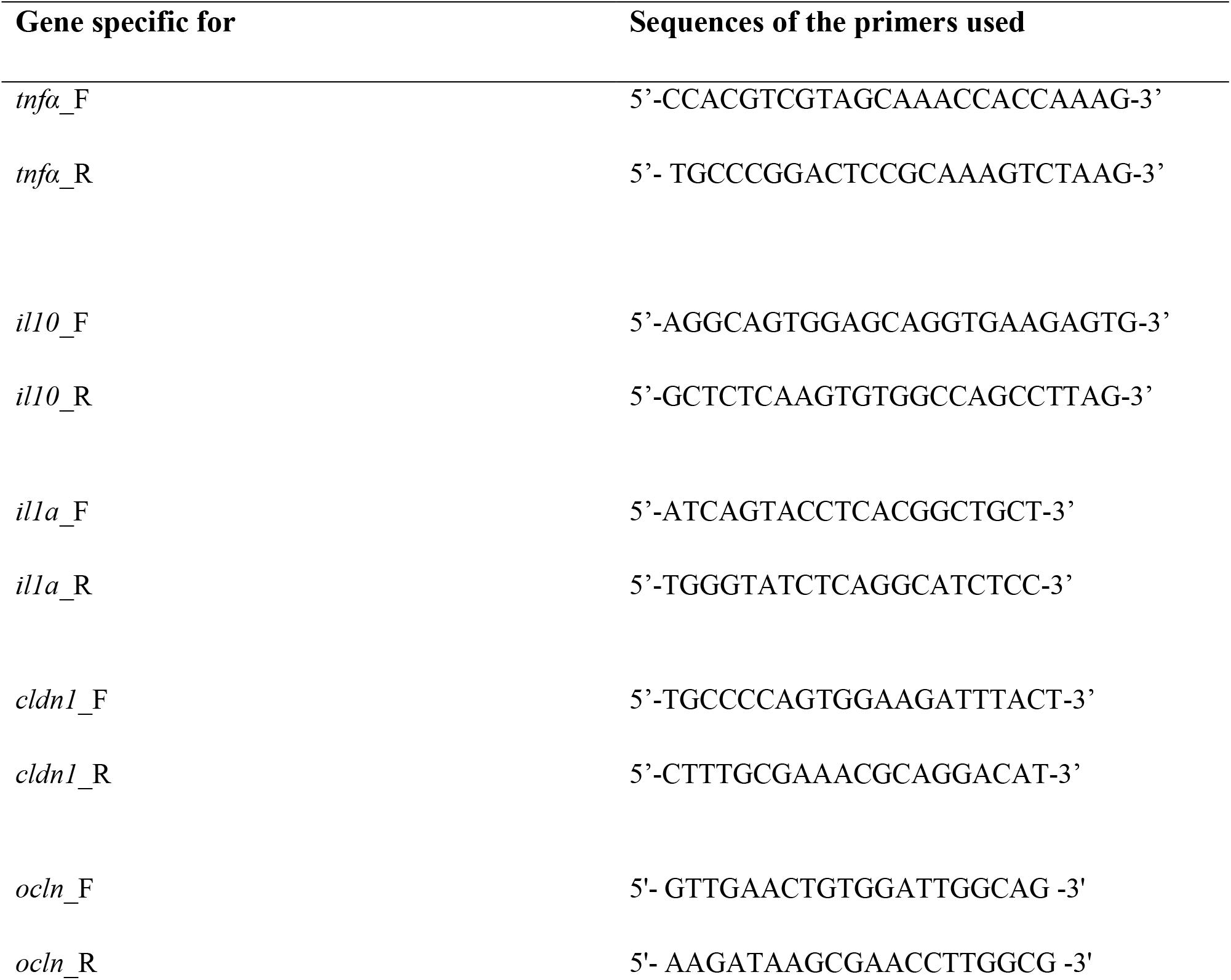
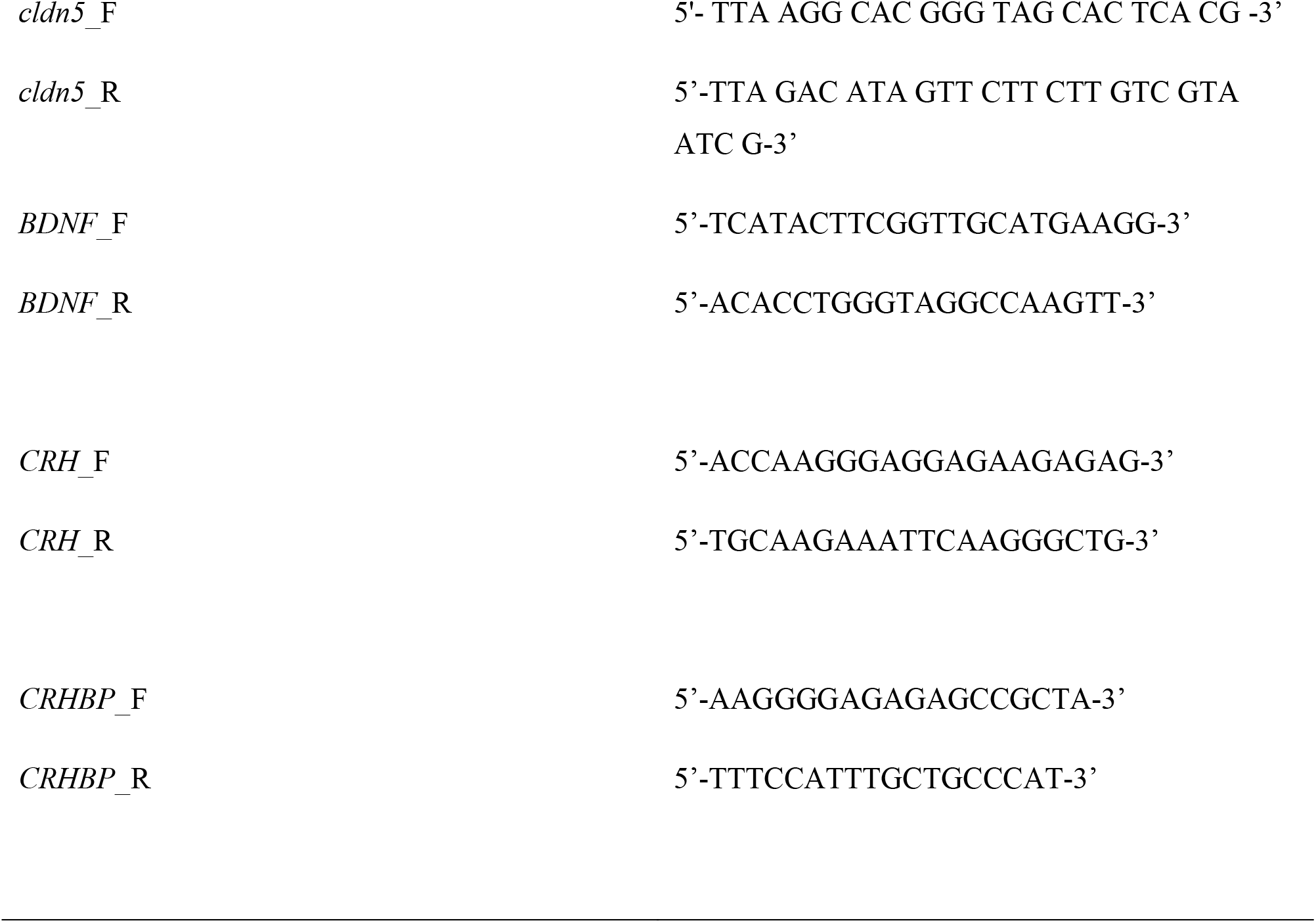
Sequences of forward (_F) and reverse (_R) primers for PCR studies to confirm presence and expression level of various genes used in this study.

BDNF has a significant relationship with the stress level of the host. An earlier report showed that BDNF level decreases and CRH level increases in anxiety patients than normal individuals (47). Dysbiosis of gut microbiota caused changes in BDNF and CRH levels in the host (48, 62). In the current study, we found less BDNF and higher CRH in the brain of vancomycin treated mice, which was justified by the higher anxiety and depressive behavior of vancomycin-treated mice compared to control mice in EPM, FST and OF test. Increased pathogenic Proteobacteria and decreased beneficial microbes during vancomycin treatment caused stress in mice which modulated BDNF, CRH and CRHBP levels in the brain. The expression pattern of these stress related genes are proposed to be due to the alteration pattern of gut microbes hence the behavior of mice. Increased Proteobacteria level on the fourth day and Verrucomicrobia level on the sixth day following vancomycin treatment showed two opposite effects on the expression of stress related genes. On the 15^th^ day of restoration, BALB/c mice showed more similar expression of stress genes with its respective control groups compared to C57BL/6 mice. The results and the correlation between changes in the stress associated genes and specific gut microbiota perhaps indicated a plausible causal relation and regulation for both strains of mice.

Proteobacteria phylum contains mostly gram-negative pathogenic bacteria to contribute LPS to bind to the TLR receptor of the gut and activates the expression of pro-inflammatory cytokines (63). Firmicutes, specifically Clostridium group present in the gut produces short-chain fatty acid (64) and these SCFA in the gut suppresses the LPS and pro-inflammatory cytokines and enhances the secretion of the anti-inflammatory cytokines (65, 66). In the current study, increased Proteobacteria and decreased Firmicutes enhanced the inflammation and permeability of gut and brain in mice, while increased Verrucomicrobia alleviated these effects during vancomycin treatment. All the major changes that happened in the host during the perturbation period became normal with the successful restoration of gut microbes.

In summary, gut microbiota perturbation and restoration followed some specific patterns following vancomycin treatment. Alteration in the abundance of a few specific groups of gut microbes mostly regulated the behavior and immune system of mice. The alteration patterns (both during perturbation and restoration period) of gut microbes were time dependent and significantly varied between BALB/c and C57BL/6 mice.

## Acknowledgement

This research received no specific grant from any funding agency in the public, commercial, or not-for-profit sectors.

## Conflict of Interest

The authors declare that there is no conflict of interest.

## Funding & payment

The current work (necessary resources to perform the experiment and the infra-structure for the laboratory) was supported by the parent institute National Institute of Science Education and Research. The current work was not supported through any extra-mural funding except the Ph.D. fellowship to PR by the Council of Scientific and Industrial Research (CSIR), Govt. of India, India. The current authors have no support to pay for the open-access or article processing fees to publish this research article.

